# GeneBag: training a cell foundation model for broad-spectrum cancer diagnosis and prognosis with bulk RNA-seq data

**DOI:** 10.1101/2024.06.27.601098

**Authors:** Yuhu Liang, Dan Li, Aguix Guohua Xu, Yan Shao, Kun Tang

## Abstract

Numerous Pre-trained cell foundation models (CFM) have been crafted to encapsulate the comprehensive gene-gene interaction network within cells, leveraging extensive single-cell sequencing data. These models have shown promise in various cell biology applications, including cell type annotation, perturbation inference, and cell state embedding, etc. However, their clinical utility, particularly in cancer diagnosis and prognosis, remains an open question. We introduce the GeneBag model, a novel CFM that represents a cell as “a bag of unordered genes” with continuous expression values and a full-length gene list. Pre-trained on single-cell data and fine-tuned on bulk RNA-seq datasets, GeneBag achieves superior performance across cancer diagnosis and prognosis scenarios. In a zero-shot learning setting, GeneBag can classify cancer and non-cancer tissues with approximately 96.2% accuracy. With fine-tuning, it can annotate 40 different types of cancers and corresponding normal biopsies with an overall accuracy of ∼97.2%. It notably excels in classifying challenging cancers such as bladder (93%) and stomach (90%). Furthermore, GeneBag is capable of cancer staging with 68.5% accuracy and 5-year survival prediction with an AUC of ∼80.4%. This study marks the first to demonstrate the potential of CFMs in RNA-based cancer diagnostics and prognostics, indicating a promising avenue for AI-assisted molecular diagnosis.

## Introduction

Traditionally, systems biology has aimed to fully understand the entirety of multi-omics data and to map the intricate relationships between countless biomolecules. The aspiration is that a complete analysis of biological systems will facilitate an in-depth comprehension of the underlying mechanisms, thereby significantly advancing disease diagnostics and the innovation of pharmaceuticals and therapeutics (G. Li et al. 2022; X. Wang et al. 2021; Kitano 2002). However, efforts in large-scale systems biology research were significantly impeded by the constraints of existing computational algorithms and analytical techniques, which struggle to keep pace with the intricate complexity and vast scales inherent in living organisms. Consequently, the findings from these studies often remain descriptive, and their predictions usually diverge significantly from in vivo conditions, limiting their practical applicability in real-world scenarios (Gomez-Ramirez and Sanz 2013; Bartocci and Lió 2016; Atta and Fan 2021).

Recent artificial intelligence research has seen remarkable progress, particularly with the advent of foundation models based on transformer architecture. Models like BERT, GPT employ self-attention to weigh and distill the complex interplays among thousands of words within the sentence contexts (Floridi and Chiriatti 2020; Devlin et al. 2018). Once trained on a vast corpora of natural language, the models have grasped the subtlest “rules of speaking,” owing to the deep neural networks’ ability to fit all conceivable functional relationships. This development is highly inspirational for systems biology, where biomolecules, akin to words, engage in intricate communications. The potential to learn comprehensive molecular interactome within a foundation model framework suggests that downstream biological inferences could be conducted, analogous to the fine-tuning or prompting techniques employed in large language models.

Indeed, a multitude of cell foundation models (CFMs) have recently emerged, trained on extensive single-cell RNA sequencing (scRNA-seq) datasets. Models such as scBERT (F. Yang et al. 2022), Geneformer (Theodoris et al. 2023), GeneCompass (X. Yang et al. 2023), and scFoundation (Hao et al. 2023) are primarily based on the BERT-like bidirectional transformer architecture, while scGPT adopts a GPT-like generative transformer approach (Cui et al. 2024). The majority of these CFMs consider gene order in their input processing (e.g., Geneformer, GeneCompass, scGPT, UCE), but scBERT and scFoundation stand out by being agnostic to gene order, utilizing gene ID embeddings instead of positional embeddings. Furthermore, while many models necessitate the discretization of continuous expression values, GeneCompass and scFoundation opt for embeddings of continuous values, which more accurately reflect the raw scRNA-seq data. These CFMs have demonstrated a variety of applications in downstream tasks, including cell type annotation, single-cell perturbation inference, target gene prediction, and drug response forecasting, showcasing their versatility and potential in advancing single-cell biology research.

However, the potential CFMs in clinical scenarios, particularly in cancer diagnosis and prognosis, has not yet been fully explored. Theoretically CFMs could apply the intergenic interaction patterns learned from one data modality, such as scRNA-seq, to identify diseases across various other data modalities, such as bulk RNA sequencing data (bulk RNA-seq), using transfer learning. Traditional bulk RNA-seq has been a cornerstone in cancer research, with different RNA components showing distinct signatures associated with cancer development (H. Wang et al. 2022). Protein-coding mRNAs have shown aberrant expression profiles across various cancers, prompting the proposal of mRNA expression profiling panels for diagnostic purposes (Bareche et al. 2018; Golub et al. 1999; Ma et al. 2020; Teresa Agulló-Ortuño, López-Ríos, and Paz-Ares 2010). Circular RNAs (circRNAs), noted for their differential expression in different cancer types (Shang et al. 2019; Feng et al. 2022; Xia et al. 2018), are considered promising biomarkers (J. Li et al. 2020; Meng et al. 2017; S. Wang et al. 2021; Wen, Zhou, and Gu 2020; H. Zhang et al. 2017). Other RNA species, such as long noncoding RNAs (lncRNAs) and micro-RNAs (miRNAs), have also been implicated in signaling specific cancers (Shen 2020; Hua, Chen, and He 2019; Loewen et al. 2014; Y. Wang et al. 2015; Xu et al. 2020; Zhan et al. 2020; Giulietti et al. 2018; Wu et al. 2021; Iorio and Croce 2012; Lu et al. 2005; Valihrach, Androvic, and Kubista 2020; Verduci et al. 2019). However, the traditional focus has been on identifying a limited set of biomarkers representing only a fraction of cancer subtypes. These marker-gene-based approaches do not enable a comprehensive assessment of the entire spectrum of cancers. The heterogeneous nature of oncogenesis further complicates the representation of all cancer subtypes by biomarker panels. Recently, there has been a shift towards employing machine learning or deep learning techniques to analyze complete omics data, aiming to provide a more profound understanding of disease mechanisms (Nicora et al. 2020) and to improve tumor classification and prognosis (L. Zhang et al. 2018; Bostanci et al. 2023). However, these applications are currently confined to specific cancer types and rely on end-to-end training with limited datasets.

In this study, we explore the utilization of CFM for the first time in the context of cancer diagnostics and prognostics, leveraging bulk RNA-seq data. We constructed a novel foundation model, the GeneBag, underpinned by a transformer encoder architecture, capable of concurrently processing all genes with an assumption of random gene order and continuous expression values. Initially trained on an extensive dataset of single-cell transcriptomics, the model was subsequently retrained using bulk RNA-seq data. We then evaluated its performance across a series of downstream tasks of cancer diagnosis and prognosis.

## Results

In general, cell foundation models aim to capitalize on transformer structures to discern gene-gene interactions from transcriptomic data, in a manner akin to language learning. However, there are notable distinctions between cellular transcriptomic data and language constructs. Language sentences consist of discrete word tokens, with a strict sequence critical for conveying precise meaning. In contrast, transcriptomic data typically regard genes as an unordered set, with gene expressions represented as continuous scalar values post logarithmic transformation of the raw count data (F. Yang et al. 2022). In this study, we intentionally designed the GeneBag model, a CFM that adapts to the unique characteristics of transcriptomic data, ensuring optimal alignment with biological frameworks. Like scBERT, we employed a bidirectional encoder, substituting positional embeddings with gene id embeddings (Fig. 1). The model’s indifference to the sequence of gene tokens was reinforced through iterative shuffling during both pretraining and fine-tuning stages (Methods). Distinct from scBERT, we incorporated continuous value embeddings to mirror the quantitative spectrum of gene expressions (methods). Gene expressions were processed and normalized individually within each gene across the training dataset (see Methods). Additionally, GeneBag leverages the Longformer architecture (Beltagy, Peters, and Cohan 2020) to facilitate the comprehensive analysis of the complete gene list, encompassing 17,930 genes across all datasets in this study, in a single operation (methods).

**Figure 1.**
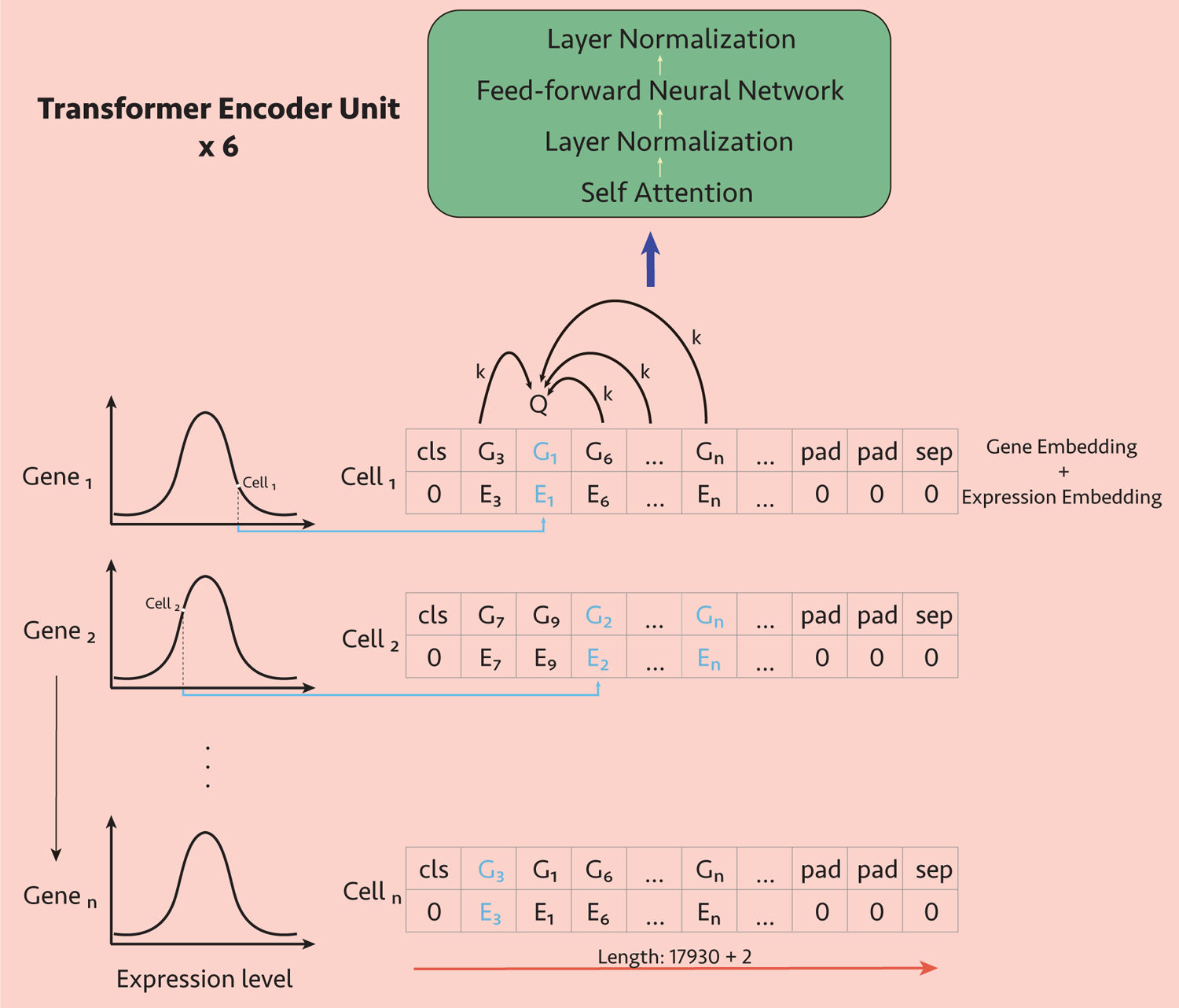
Model structure Each cell consists of a pair of gene and expression sequences. Gene IDs were tokenized, paired with the corresponding expression value embeddings to generate gene embeddings. In pre-training and fine-tuning, these embeddings, representing 17,930 genes, are fully shuffled to enhance model generalization before being fed into the encoder. The encoder consists of 6 layer transformers with multi-head attention. The output from the encoder is subsequently processed by task-specific decoder networks for fine-tune tasks.

### Pretraining and Evaluation with Single-Cell RNA Sequencing Data

GeneBag underwent initial pretraining on 1.3 million human single-cell RNA sequencing (scRNA-seq) data points from PanglaoDB (Franzén, Gan, and Björkegren 2019), employing a strategy of random masking of either gene identifiers or expression values (Fig. 1). This pre-training phase endowed the model with the capability to impute expression values for any random genes with a remarkable accuracy of 94.9% (Pearson correlation coefficient), indicative of the model’s adept learning of gene-gene interactions at the single-cell level.

Our initial evaluation of GeneBag focused on the task of single-cell annotation, utilizing the Zheng68K dataset as a benchmark (Zheng et al. 2017). This dataset, comprising blood mononuclear cells of highly similar subtypes, presents a great challenge to cell annotation methodologies (F. Yang et al. 2022). Post fine-tuning, GeneBag achieved an overall accuracy of 71.33% and an F1-score of 0.61. The cell type annotation closely mirrored the ground truth, as visualized on the T-SNE plot (Figure 2A). The confusion matrix (Figure 2B) revealed that the majority of cell types attained a high degree of accuracy, ranging from 65.95 to 96.94%, with the exception of CD4+/CD45RA+/CD25-Naïve T, which registered a lower accuracy of 40.57%. Notably, the model demonstrated a significantly elevated accuracy of 88.94% for CD4+/CD25 T Reg, surpassing the performance of preceding models(F. Yang et al. 2022). Additionally, GeneBag was successful in distinguishing between CD8+ Cytotoxic T cells and CD8+/CD45RA+ Naive Cytotoxic cells with relatively high accuracies (65.95% and 88.42% respectively). This is particularly noteworthy given that these cell types have traditionally been considered challenging to discern from one another due to their marked similarities(Zheng et al. 2017)(F. Yang et al. 2022).

**Figure 2.**
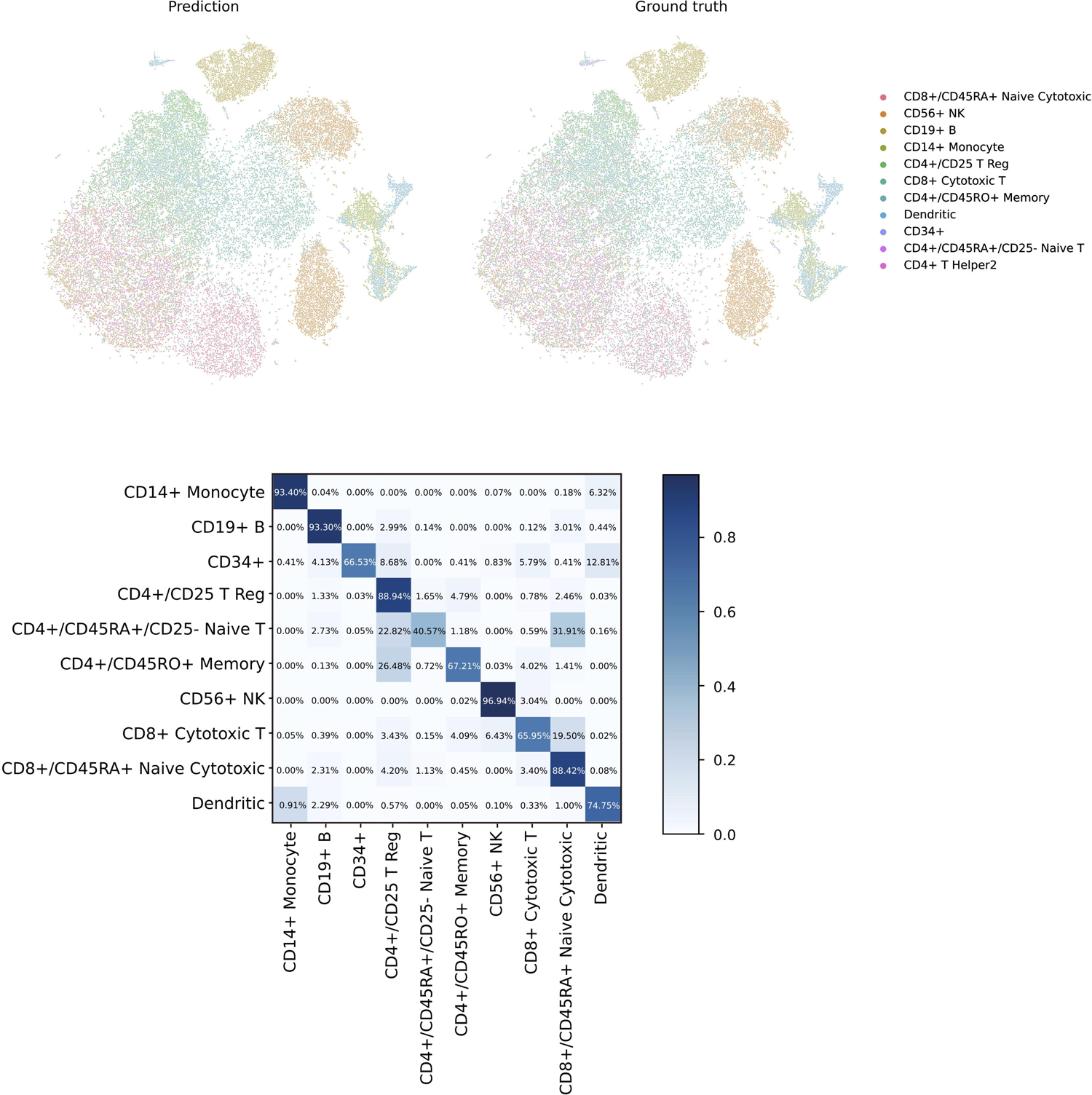
Single cell annotation. A. In the scatter plots of Zheng68K cells, X and Y positions were obtained from T-SNE analysis of raw expression. Colors represent the different cell type labels. B. Confusion matrix on the Zheng68k test cells. Columns represent ground truth and rows represent the predictions.

### Transfer Learning of the GeneBag Model on Bulk RNA-seq Data

In clinical practice, tissue-level bulk RNA sequencing is a widely utilized and well-established diagnostic method. Should the cell foundation model successfully integrate bulk RNA-seq data, it would pave the way for numerous downstream applications in diagnostics and prognostics. Self-attention based foundation models have demonstrated remarkable capabilities in multimodal tasks, particularly through transfer learning, across domains such as natural language processing and computer vision (Zhu et al. 2020; Vaswani et al. 2017). Motivated by this, we explored the application of the GeneBag model to bulk RNA-seq data through a retraining approach. We retrained GeneBag on a mixed dataset, integrating 100k single cell samples from PanglaoDB and 19k bulk RNA-seq samples from the Genotype-Tissue Expression (GTEx) project (Lonsdale et al. 2013; Wilks et al. 2021) (see Methods). For training the bulk RNA-seq data, the same gene ID embedding and expression embedding methods was used as for the scRNA-seq. The <*cls*> token was used to distinguish the datatypes of scRNA and bulk-RNA. This hybrid retraining strategy facilitated the model’s adaptation to the nuances of bulk RNA-seq data, while minimally impacting its proficiency in the single cell context, as evidenced by the overall mask prediction accuracy of 90.51%.

### Tissue and Cancer Classification utilizing bulk RNA-seq

Upon retraining the GeneBag model, we initially assessed its capacity to classify various tissues and cancer types using bulk RNA-seq data. The primary inquiry was whether the model could identify the intrinsic differences between normal and malignant tissues, potentially enabling it to differentiate between these states in previously unseen tissue types through zero-shot learning. To examine this, we selected a diverse range of ten tissue types—encompassing breast, colon, esophagus, kidney, liver, lung, prostate, stomach, thyroid, and uterus—where both normal (from GTEx) and tumorous (from TCGA (Wilks et al. 2021)) samples were available. The samples of these tissues were combined into a single training set, categorized as either normal (5,397 instances) or tumorous (6,163 instances). GeneBag underwent fine-tuning on this training set. Subsequently, the model’s efficacy was evaluated using samples from tissue types not represented in the training set. The zero-shot learning prediction accuracy (on unseen tissue types) achieved an overall accuracy of 96.17%, with an exceptionally high classification specificity of 99.76% and a sensitivity for identifying tumors of 85.53% (table 1). These results imply that GeneBag has adeptly learned to generalize the distinguishing features between normal and tumorous tissues, irrespective of the considerable variations across different tissue profiles.

**Table 1.** Statistical results of Zero-shot learning task.

|  | No. of Predicted<br>Normal Samples | No. of Predicted<br>Tumor Samples |
| --- | --- | --- |
| No. of Normal<br>Samples | 13047 | 637 |
| No. of Tumor<br>Samples | 32 | 3766 |
| Sum of<br>Prediction | 13079 | 4403 |
| % of Correct<br>Prediction | 99.76% | 85.53% |

Since most cancer types have been extensively studied, a more critical question is whether a CFM, after thorough fine-tuning, can accurately classify a comprehensive range of tissue and tumor types. To address this, we curated a broad bulk RNA-seq dataset, establishing 28 distinct normal tissue (NT) types, 7 paracancerous tissue (PT) types, and 31 tumor (TM) types. We evenly divided the samples of each type into training and testing sets with an 8:2 ratio. GeneBag was then subjected to fine-tuning on the training dataset, followed by testing on the reserved test set. The output embeddings from the classification network were visualized using UMAP (Figure 3A). It was observed that the data points of the 66 NT, PT, and TM types formed compact clusters within the same type, while being almost distinctly separated across different types (Figure 3A). The overall classification accuracy achieved was 97.23%. Upon closer examination, NT types were found to gravitate towards the upper right, distinctly separated from the TM types clustered at the lower left, with PT samples interspersed in between (Figure 3B). Notably, NT and TM from the same tissue type (in the same color-code, in Figure 3B) were distant in the UMAP, exhibiting a similar diagonal projection, contrasting with previous studies where NT and TM of the same tissue were in closer proximity than those from different tissues (Prada-Luengo et al. 2023). It is also insightful to explore the relationships among related tissues. Figure 3C illustrates that in normal tissues, the colon and small intestine are closely aligned, mirroring their anatomical proximity, yet they are notably distant from the esophagus and stomach, indicating significant developmental divergence among these clusters (Figure 3C). Interestingly, breast samples partially intermingle with adipose tissue, hinting at a shared developmental lineage (Figure 3C). In the realm of tumor tissues, three subtypes of kidney tumors: chromophobe, renal papillary cell carcinoma, and renal clear cell carcinoma, are clearly demarcated from one another and are also far removed from the kidney paracancerous tissue (Figure 3D). This illustrates GeneBag’s proficiency in classifying tumor subtypes of the same tissue origin, a highly desirable feature for clinical specimen analysis. The model’s capacity for subtype classification is further exemplified in Figure 3E, where a total of 15 diverse tumor types are largely distinguishable within a small area of the 2D UMAP. This includes the delineation of a PT of lung and two subtypes of lung cancer (LUAD and LUSC) and two of uterine cancer (UCS and UCEC). Notably, testicular germ cell tumor, cervical squamous cell carcinoma and endocervical adenocarcinoma (TGCT and CESC) are observed to be in close proximity, suggesting a shared biological signature (Figure 3E). Similarly, mesothelioma and sarcoma of soft tissue (MESO and SARC) exhibit a comparable pattern of clustering. In summary, the developmental relatedness is reflected in the classification embeddings, yet even highly similar subtypes can be effectively segregated.

**Figure 3.**
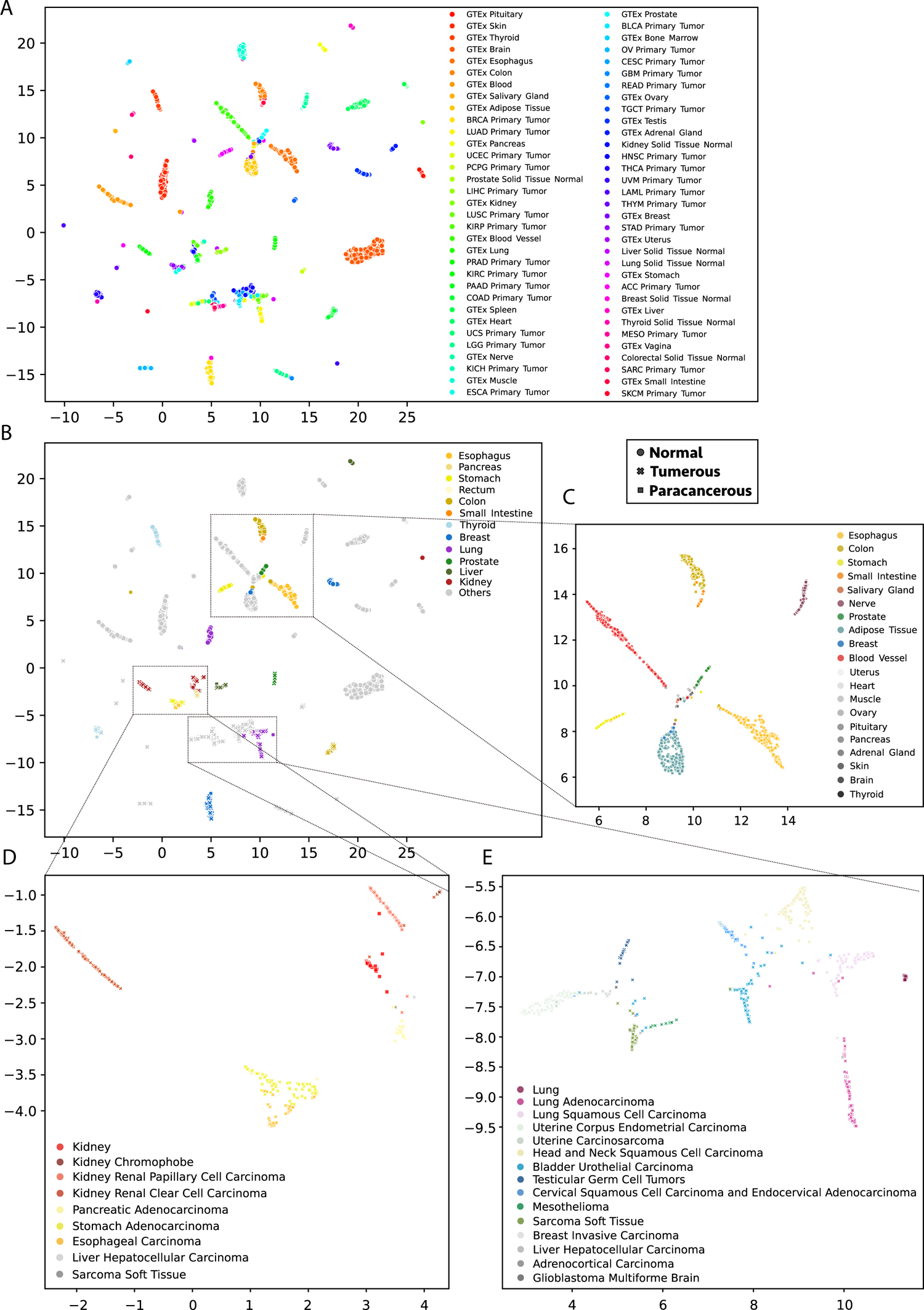
Tissue classification. A. Inference on normal and tumor samples was done with a classifier. The last layer of the classifier was used for producing UMAP. Colors represent the ground truth of the sample types. B. Same as A but with colors labeled only for 12 tissues, for which tumorous and normal types are both present. Different marker shapes represent different tissue types, with solid dots, crosses and solid squares representing normal, tumorous and paracancerous tissues respectively. C, D, E. zoom-in views of panel B.

To systematically evaluate the classification performance, we generated a confusion matrix encompassing the 40 tumor types (Figure 4A). The model’s assignments were found to align closely with the actual labels, achieving an overall accuracy of 93.62%. However, metastatic or recurrent tumors, with the exception of skin cutaneous melanoma (SKCM) metastasis which was correctly classified in 86% of cases, were predominantly categorized under their respective primary tumors, including BRCA, GBM, LGG, OV, TGCT, and THCA (Figure 4A). This observation may imply either a scarcity of distinct molecular markers for most tumor recurrences or a current limitation in GeneBag’s ability to differentiate between primary and recurrent tumors within these categories. Comparing our results with those from prior studies, our model has attained state-of-the-art (SOTA) performance across nearly all commonly studied cancer types. Particularly noteworthy is the model’s performance in classifying Bladder Urothelial Carcinoma (BLAC) and Stomach adenocarcinoma (STAD), which have been challenging to discern with bulk RNA-seq in previous studies (<0.45 for BLAC and <0.35 for STAD) (Prada-Luengo et al. 2023; Vivian et al. 2020). Our model, however, has demonstrated the capacity to classify these with high precision, attaining accuracies of 0.93 for BLAC and 0.90 for STAD. This advancement highlights the significant potential of GeneBag in enhancing the classification of historically difficult-to-identify tumor types.

**Figure 4.**
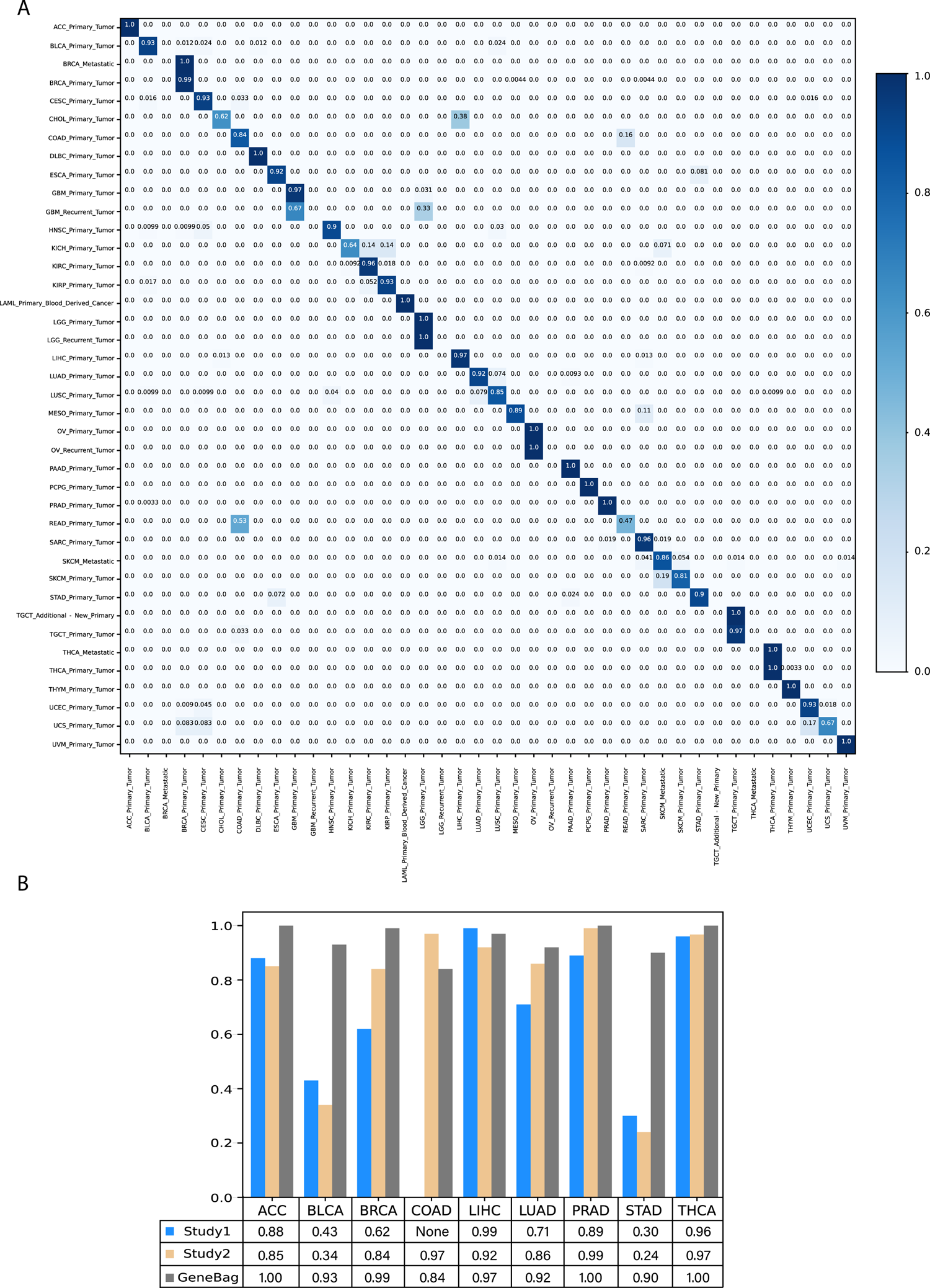
Tumor subtyping. A. Confusion matrix of tumor subtypes. Columns represent ground truth and rows represent the predictions. B. Comparison between our model and previous methods for tumor identification.

### Stage and Survival Prediction Utilizing Bulk RNA-Seq

Beyond facilitating cancer diagnosis, there is a significant interest in leveraging bulk RNA-seq data to predict cancer stages and survival outcomes. Accurate staging is essential for assessing disease severity and formulating treatment plans. However, the subjectivity inherent in cancer staging can lead to variability in results, influenced by factors such as observer differences, tumor characteristics, and diagnostic methodologies. It has been reported that staging concordance rates can range widely, from 58% to 86% (Dolly et al. 2020) (Gwon et al. 2024; Plichta et al. 2019). In our study, we explored the potential of using bulk RNA-seq from cancer biopsies to predict cancer staging across various cancer types. We included all tumor samples with available staging information and utilized paracancerous samples as a baseline stage 0 for our fine-tuning experiments. As depicted in Figure 5A, a clear correlation was observed between the predicted and actual stages. The overall Pearson correlation coefficient, measuring the relationship between predicted and labeled stages, is 0.685. This correlation is notably high, especially considering the inherent variability in staging assessments and the diverse range of cancer types included in this analysis. This result underscores the promise of the foundation model in objective and precise cancer stage estimation through the analysis of bulk RNA-seq data.

**Figure 5.**
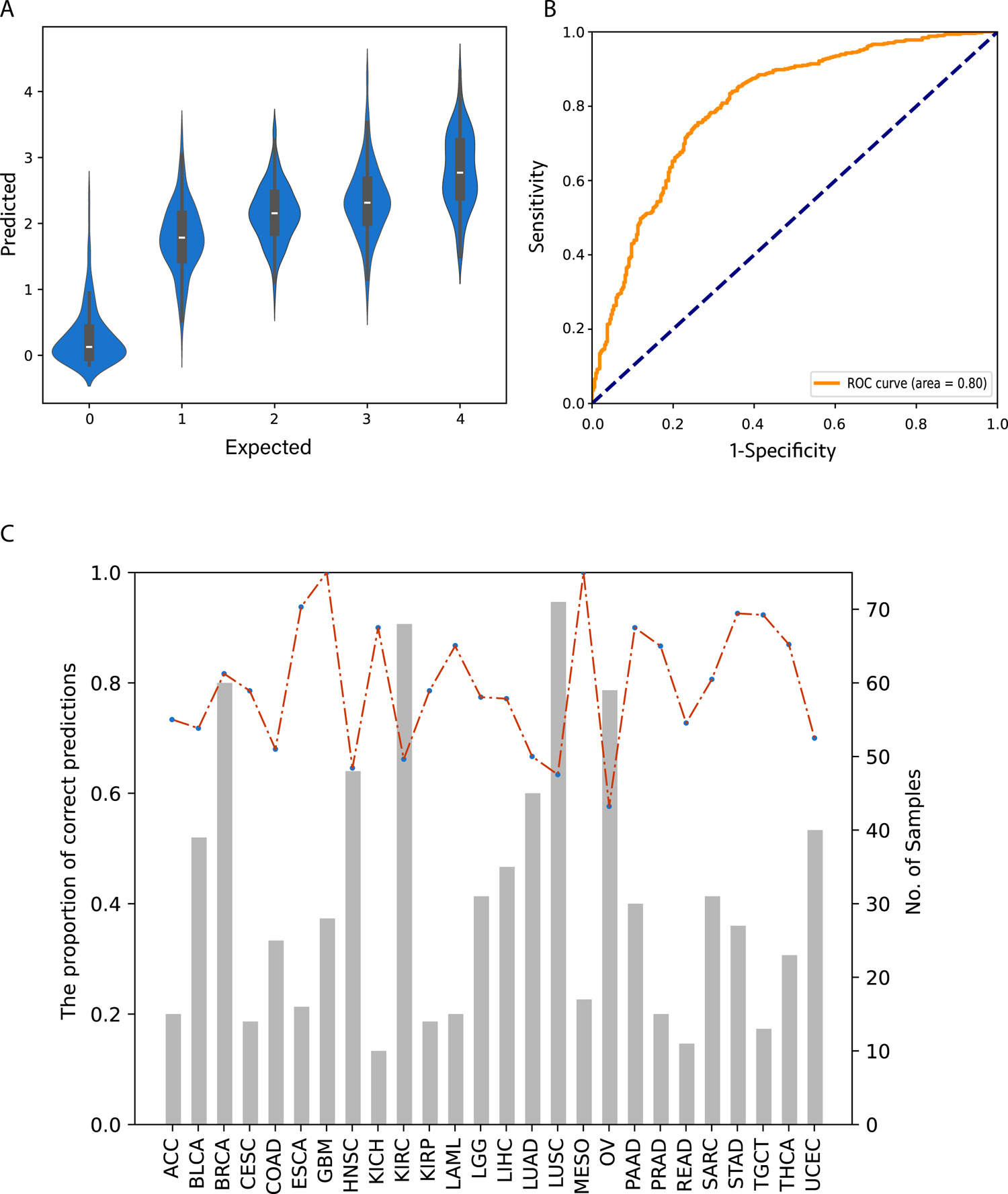
Tumor staging and survival prediction. A. Violin plot of tumor stage, X axis shows the tumor stage collected from TCGA metadata. The Y axis presents the predicted tumor stages of the GeneBag model. B. AUC plot of survival prediction. C. Bar-plot shows the sample number of each tumor type, dash-line shows the predicted accuracy of each tumor type.

Further extending our investigation, we assessed the model’s capability to predict 5-year survival outcomes using TCGA samples. The samples were categorized based on two conditions: 1, those who succumbed to the disease within 5 years post-initial cancer diagnosis (<5 years survival); 2, those who survived beyond 5 years from diagnosis or were confirmed alive after a minimum of 5 years follow-up (>5 years survival). The fine-tuned model achieved an overall accuracy of 74.94% in predicting survival rates. It is observed that the model exhibits a slightly higher accuracy in forecasting the >5 years survival group, with 81.4% of the predictions correctly identifying individuals who survived over 5 years. Conversely, the prediction for the <5 years survival group was comparatively less precise, with 65.9% of the predictions correctly aligning with those who did not survive beyond 5 years. Adjusting the prediction threshold, defined by the ratio of softmax outputs from the binary classifier network, reveals a corresponding variation in specificity and sensitivity. The model demonstrated an overall AUC (Area Under the Curve) value of 80.42% for the ROC (Receiver Operating Characteristic) curve, as illustrated in Figure 5B. It is important to note that the accuracy of survival rate predictions varied significantly across different cancer types. Notably, Esophageal Adenocarcinoma (ESCA, 93.75%), Glioblastoma Multiforme (GBM, 100%), Kidney Clear Cell Carcinoma (KICH, 90%), Mesothelioma (MESO, 100%), Pancreatic Adenocarcinoma (PAAD, 90%), Stomach Adenocarcinoma (STAD, 92.59%), and Testicular Germ Cell Tumors (TGCT, 92.31%) showed accuracies exceeding 90% (Figure 5C). In contrast, certain cancer types such as Head and Neck Squamous Cell Carcinoma (HNSC, 64.58%), Kidney Renal Clear Cell Carcinoma (KIRC, 66.18%), Lung Squamous Cell Carcinoma (LUSC, 66.38%), and Ovarian Cancer (OV, 57.63%) were found to have comparatively lower prediction accuracies.

## Discussion

In the realm of leveraging AI technologies for large-scale omics data analysis, previous studies have predominantly employed end-to-end training network approaches. These methods, while valuable, encounter limitations due to constraints such as limited labeled data and a narrow scope of application. This study, to our knowledge, is the first to make use of a Cell Foundation Model for addressing pivotal questions in cancer diagnosis and prognosis using bulk RNA-seq data. Despite having a modest number of parameters, GeneBag has demonstrated exceptional capabilities in classifying a diverse array of tissue and cancer types with high accuracy, as well as in forecasting cancer stages and 5-year survival rates. Larger models trained on more expansive single-cell data corpora would certainly further amplify performance. Moreover, with fine-tuning, foundation models may be able to address more complex questions, such as forecasting the effectiveness of diverse treatments or customizing personalized therapeutic strategies based on the analyses of bulk RNA-seq data. To accomplish this, it is essential to have cohort data that comprehensively records treatments and their associated outcomes, enabling CFMs to discern the relationship between therapeutic interventions and RNA profiles.

Traditionally, pathological testing through standard biopsy involves sectioning, staining, and microscopic examination of tissue by pathologists to ascertain cancer type and grade, a process prone to inter-observer variability. Furthermore, subtle distinctions between tumor types might not be apparent through morphological changes alone. Conversely, a CFM-based approach with biopsy bulk RNA-seq offers high-throughput, objective screening across a wide range of cancer types, and potentially with greater sensitivity and specificity due to the utilization of comprehensive transcriptomic data. Additionally, CFMs could enable the inclusion of new inferences in a single examination, such as prognostic assessments and precision medicine strategies. Ultimately, it is anticipated that such novel, foundation model-driven and RNA profiling based diagnostic approaches will complement traditional pathology, offering supplementary information to bolster clinical decision-making. However, the integration of CFMs into clinical practice, like any AI tool, necessitates stringent validation and regulatory approval to ensure safety and efficacy.

## Methods

### Overview

The GeneBag Model is a modified BERT (Bidirectional Encoder Representations from Transformers) model. It processes input sequences from single-cell or bulk tissue RNA-Seq data through an encoder with multi-head attention mechanism. The encoder output is then processed differently during the pre-training and fine-tuning stage. In pre-training (and also in retraining), it performs a fill-mask task, using a decoder to predict expression values for masked genes. This trains both the encoder and decoder. During fine-tuning, a classifier is trained by applying it to the entire encoder output, adapting the model for specific prediction about each sample.

While this process shares similarities with other language models used in gene expression analysis, GeneBag introduces three key innovations. 1. GeneBag doesn’t rely on fixed gene positions or position encoding, only requiring alignment between gene tokens and expression values. 2. Unlike binned expression, GeneBag directly encodes and decodes raw expression data, preserving the full granularity of the biological information. 3. By incorporating the Longformer attention module, GeneBag can process input sequences containing full gene lists.

### Input Embedding

We collected a huge RNA-Seq dataset and organized it into paired sequences. In the dataset, each sample consists of a gene ID sequence and its corresponding expression value sequence. We incorporated special tokens (*<CLS>, <PAD>,* and <*sep*>) at the beginning and end of sequences, similar to other language models. All gene IDs and special tokens were tokenized as integers using a common vocabulary. Unlike other models, genes don’t have fixed positions in our sequences. However, gene ID tokens remain aligned with their corresponding expression values in the paired sequences, maintaining crucial gene-expression relationships. Gene ID tokens were converted to gene embeddings of dimension *d_model* using PyTorch torch.nn.embedding. Expression values were directly encoded into expression embeddings of the same dimension *d_model* using a series of cosine and sine functions. This deterministic encoding allows for precise representation of continuous data. The gene and expression embeddings were then added together to form the input (1) for the Longformer encoder.

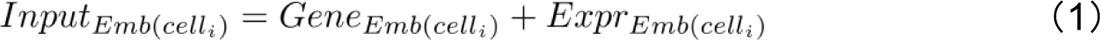

### Encoder with attention mechanism

Usually the input embeddings were processed with a transformer encoder. However, standard self-attention struggles with very long sequences due to its quadratic complexity. This is problematic for transcriptome data, which can exceed 15,000 genes. To solve this, we implement Longformer attention, which uses a sparse attention pattern. This allows the model to handle longer input sequences efficiently and potentially capture broader biological contexts and interactions. Our sequence length is set to 17,932. Our Longformer encoder includes six attention heads and six encoder layers. The model’s dimension (*d_model*) is set to 42.

### Pre-training of GeneBag Model Based on Single Cell RNA-seq Data

The pre-training data consists of approximately 1.3 million single-cell RNA-Seq sequences. Each cell sequence was firstly processed with masking, where 15% of the gene expression values in the non-padding regions are randomly replaced. These masked sequences are then processed through input embedding and the Longformer encoder. The encoder output for each masked gene is fed into a multi-layer perceptron to predict the expression value. The predicted expressions are compared against the actual values using Mean Squared Error (MSE) as the loss function. Prediction accuracy is evaluated with the Pearson correlation coefficient. Other configurations include a batch size of 32, a learning rate of 1e-4, and the Adam optimizer.

### Cell Type Annotation

We benchmarked the GeneBag model’s performance in cell type annotation using the Zheng68K dataset. In this dataset, the occurrence of some cell types is sparse, leading to an imbalanced distribution. To address this, we set a threshold of 6,185 cells per cell type. For types exceeding this threshold, we randomly select a subset to meet this limit. Cell types with fewer than 6,185 cells are fully included, creating a more balanced dataset. This process results in a dataset with a total of 40,828 cells. This was further divided as a train and test dataset with 8:2 ratio. A classifier with a single convolutional layer followed by three multilayer perceptrons (MLPs) was used to predict the cell type from the entire encoder output. The loss function is CrossEntropy.

### Re-training of GeneBag Model on Bulk RNA-seq

Re-training is similar to pre-training while including the bulk tissue data. We used the entire GTEx data (19,081 samples) and randomly sampled 100,000 cells from the pre-training dataset, labeled as “bulk” and “scRNA”, respectively. We merged them to create a balanced dataset. We further divided it into training/test/validation subsets in an 8/1/1 ratio. The <*cls*> token was used to learn the data type (bulk tissue or single cell).

### Zero-shot Tumor Recognition

To assess the GeneBag model’s capacity to accurately identify tumor samples among previously unseen tissue types, we selected ten tissues as training set in this fine-tune task. These 10 tissues were chosen based on two criteria: (a.) their presence in both the GTEx and TCGA databases; (b.) the availability of at least 50 samples from the GTEx and TCGA-tumor datasets, as well as at least 10 samples from the TCGA-normal datasets. For testing, we included 13,684 normal and 3,798 primary tumor samples from other tissue types. Similar to cell type annotation, we used the same classifier and CrossEntropy loss function to distinguish normal and tumor samples.

### Tissue and Cancer Type Classification

The whole dataset was composed of three distinct categories: normal tissues from GTEx, tumor tissues from TCGA-tumor and paracancerous tissues from TCGA-normal. We included only types of sample size exceeding 50. We evenly divided these samples into training and testing sets, comprising 23,654 and 5,945 samples, respectively. For confusion table analysis of tumor subtypes, 40 tumor types were analyzed. Tumor types with 5 or fewer samples were excluded. The total samples were evenly divided within each tumor type according to a 8:2 ratio, resulting in a training set of 7,987 and a test set of 2,525 samples. The same classifier network was used except the number of output neurons were matched to the number of types.

### Tumor Staging

For tumor staging, we focused solely on primary tumor samples for stage prediction purposes. Stage labels i to iv were used, and subtypes such as ia, ib, were merged into the major type.

Samples using non-standard labels were excluded for clarity. Solid normal tissues, which are paracancerous in TCGA, were labeled as stage 0. The final dataset comprised a total of 7,611 samples. We assumed the cancer stages as continuous values. The decoder to predict stages was similar to the cell type classifier, except the final layer of the MLP goes to a single neuron to estimate the tumor stage, and MSE loss was used.

### Survival Prediction

In the TCGA dataset, we identified 2,559 samples who did not survive beyond five years and 1,590 individuals who lived longer than five years. For the binary classification task, we applied the same classifier architecture as mentioned before.

### Datasets Panglao dataset

The Single Cell RNA-seq (scRNA) gene expression dataset was from the Panglao dataset (https://panglaodb.se/). This dataset contains 1,16,580 cells with 77 tissues, which were collected from 209 single cell datasets. In our study, only the gene IDs and expression values from the Panglao dataset were used..

### Zheng68K dataset

Zheng68K (Zheng et al. 2017) is a well annotated PBMC dataset generally used for cell type annotation evaluation, and it consists of 11 cell types, totaling 68,450 cells. Among them, there are 20,757 CD8+ Cytotoxic T cells, 16,645 CD8+/CD45RA+ Naive Cytotoxic cells, 8,775 CD56+ NK cells, 6,185 CD4+/CD25 T Reg cells, 5,877 CD19+ B cells, 3,059 CD4+/CD45RO+ Memory cells, 2,847 CD14+ Monocyte cells, 2,095 Dendritic cells, 1,871 CD4+/CD45RA+/CD25-Naive T cells, 242 CD34+ cells and 97 CD4+ T Helper2 cells, accounting for 30.3%, 24.3%, 12.8%, 9.0%, 8.6%, 4.5%, 4.2%, 3.1%, 2.7%, 0.4%, and 0.1%, respectively.

### Genotype-Tissue Expression (GTEx)

GTEx is a public database containing transcriptome gene expression data from large scale human normal tissues. We downloaded the raw gene counts from Recount3 (Wilks et al. 2021) platform via the build-in R packages. The total data of 31 tissue types with 19,081 samples were used in our study.

### The Cancer Genome Atlas Program (TCGA)

The TCGA database is composed of RNA sequencing data of tumor biopsies, with a total of 11,348 samples. As above, the raw gene counts and meta-data files both were obtained from Recount3 (Wilks et al. 2021).

### Data pre-processing

Raw expression values are all read counts. For GTEx data, we removed low-expression genes and non-protein-coding genes. This results in a list of 16,874 genes. The same genes were selected for TCGA data. Log transformation was done only for GTEx and TCGA data, because the single cell data was already processed. Lastly, we performed a gene-wised normalization by calculating z-scores for all datasets (2).

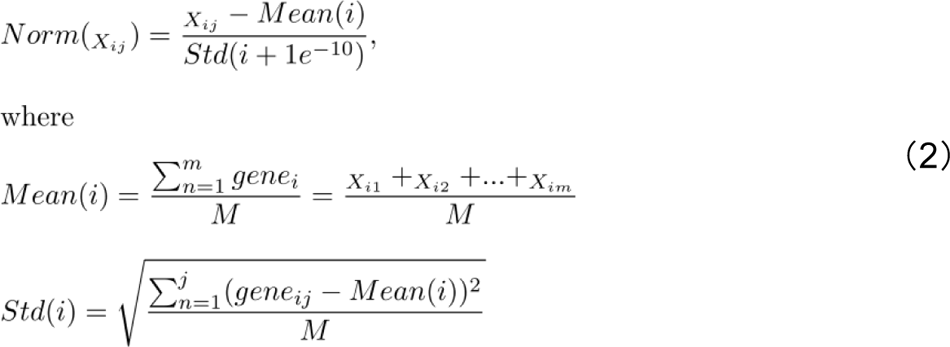

## Code availability

The code for the GeneBag model’s pre-training and subsequent fine-tuning tasks are available on Github (https://github.com/transomics/GeneBag).

## Acknowledgement

We would like to extend our sincere thanks to Zhejiang Lab. The lab provided computational resources for our model training. A special thanks goes to Prof. Xingxu Huang for his advice on our study. KT received support from Zhejiang Lab self-designed project “K2023KG0AC03”.

